# Massive lateral gene transfer under strain coexistence in the gut

**DOI:** 10.1101/2023.09.25.559333

**Authors:** N. Frazão, E. Seixas, H.C. Barreto, M. Mischler, D. Güleresi, I. Gordo

**Author notes:** These authors contributed equally (N. Frazão, E. Seixas, H.C. Barreto).

## Abstract

Mammals are colonized by multiple strains of *Escherichia coli*, yet how such strain coexistence affects their tempo and mode of evolution is poorly understood. Here, by following the colonization of two phylogenetic distinct strains of *E. coli* in the mouse gut, we find a strain-specific mode of evolution and a remarkable level of gene transfer between strains. In the same host, despite accumulating mutations at the same rate, one strain evolves by diversifying selection and the other by directional selection, and a rich dynamics of bacteriophage and plasmid transfer is found. Our results provide support for an important role of lateral transduction in the mammalian gut.

## Main

*Escherichia coli* is a highly diverse and versatile species, being both a commensal of humans and other mammals^1^, but also implicated in several intra and extra-intestinal diseases^2^. *E. coli* polymorphism for gene number and mobile genetic elements contributes to its extensive phenotypic diversity as a commensal and a pathogen^3^. A healthy human can be colonized by one dominant strain or, more commonly, by several strains^4^. Turnover of *E. coli* strains is also typically observed in the human gut, but the causes of such population structure dynamics are still not well understood. Transmission across human hosts, between animals and humans, and from the environment, is thought to be a contributor to residence times of the strains in the human gut, but not the only one^5^. Evolution by mutation and horizontal gene transfer, *via* conjugation and transduction, is pervasive in *E. coli,* and both processes are known to contribute to within-host adaptation^6–8^. Thus, the speed of evolutionary adaptation of a strain could be an important contributor to its residence time^9^. However, the extent to which rates of adaptation vary across *E. coli* strains coexisting in the same host is poorly known. Here, we take an experimental evolution approach to address this knowledge gap.

We report an *in vivo* evolution experiment where the evolution of two strains of *E. coli* is followed in real time during ∼1600 generations (3 months). We colonized mice after 7 days of antibiotic treatment, which allows for the successful colonization by *E. coli* of the mouse gut even under a complex microbiota species composition^10^ (**Fig. 1**). One strain belongs to phylogenetic group B1, which is a common resident of the mouse gut^11^. Previous studies^10,12^ indicate that it is also a good donor of Mobile Genetic Elements (MGEs) (donor strain). The other strain belongs to phylogenetic group A, is a well characterized strain and a good recipient of MGEs (recipient strain). Each strain is labelled with two neutral fluorescent markers that help to track the emergence of adaptive events during colonization. Mice were co-colonized with the two strains (n=9) or mono-colonized with each of the strains (n=3) (**Supplementary Fig. 1**). In the co-colonization we expect events of horizontal gene transfer (HGT) to be common, either driven by bacteriophages or by conjugative plasmids^10^. This additional mechanism of evolution should create new selective pressures beyond those occurring in the mono-colonization, where only mutation events are expected to fuel evolutionary change and could lead to alterations in the mode of evolution of each strain.

**Fig. 1:**
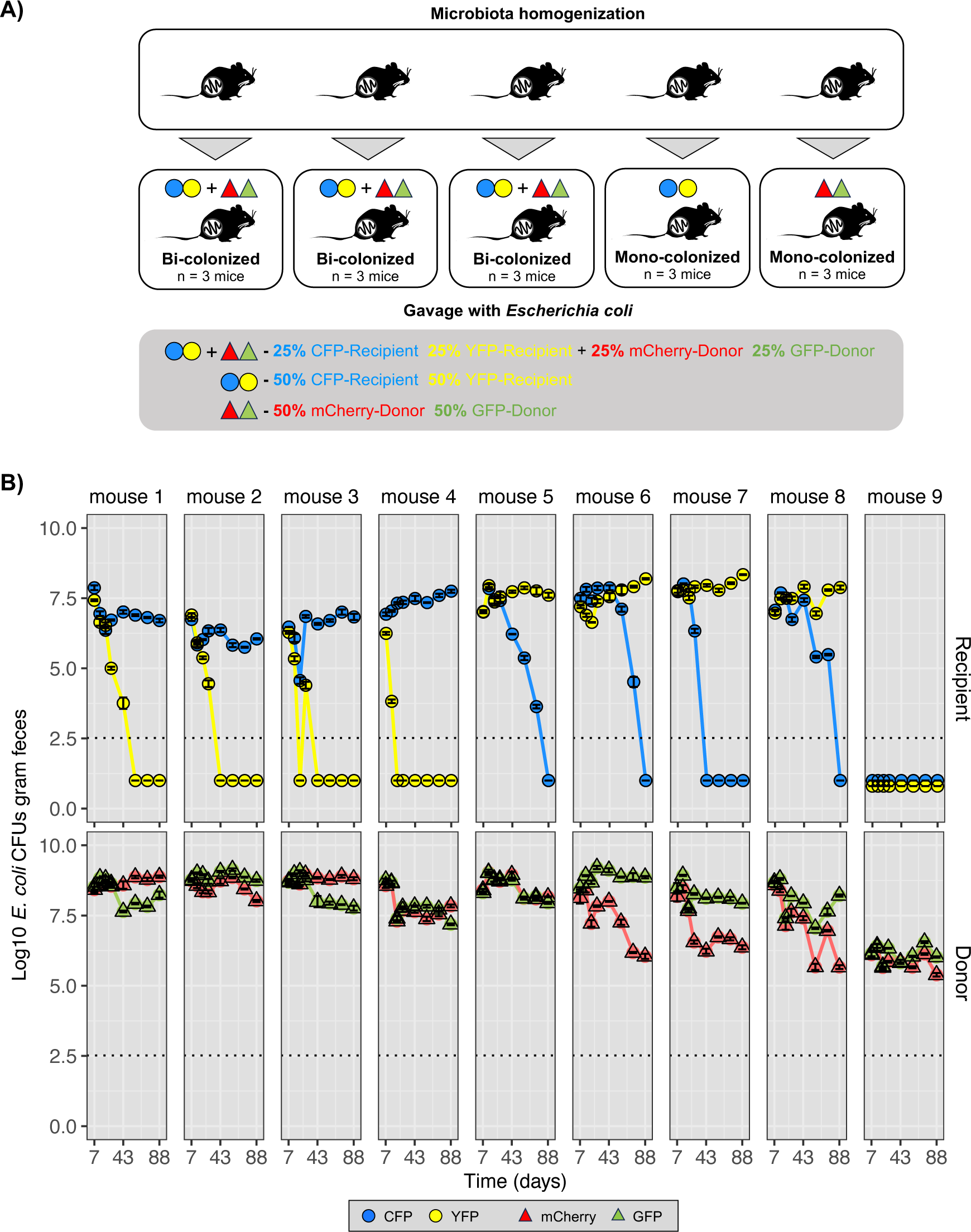
Experimental design and distinct modes of evolution between strains. **(A)** Before gut colonization, three cohorts of five mice (n = 3×5) were co-housed for 10 days to achieve microbiota homogenization: the first day without antibiotic treatment, followed by 7 days of streptomycin treatment and 2 days without antibiotics in the water to clear the gut of any streptomycin traces^23^. In each cohort, three mice were bi-colonized with donor (circles) and recipient (triangles) *E. coli* (1:1), one was mono-colonized with the donor strain and another with the recipient strain, to follow evolution in the mouse gut (see **Supplementary Fig. 1** for more details). **(B)** Time series of the abundances of the donor and recipient *E. coli* lineages carrying one of 4 fluorescent markers (CFP, YFP, mCherry and sfGFP). Dotted line represents the limit of detection for *E. coli*. Error bars represent the standard error of the mean (2*S.E.M).

Temporal series data of the abundance of each strain shows that they can coexist for at least 3 months in the mouse gut (coexistence observed in 8 out of 9 mice, **Fig. 1B** and **Supplementary Table 1**). Under coexistence, the absolute abundance of the donor strain in the feces was significantly higher than that of the recipient (mean Log_10_CFUs of donor: 8.5 (±0.4, S.E.M) g^-1^ of feces and mean Log_10_CFUs of recipient 7.4 (±0.3, S.E.M) g^-1^ of feces, Linear mixed-effects model fit by maximum likelihood, *P* = 0.0074, **Supplementary Fig. 2**). Assuming a simple evolutionary model where adaptive mutations are unconditionally beneficial and their effects are similar between strains, we would expect the rate of evolution to be higher in the donor strain, given its higher population size^13^. This should then lead to a faster rate of loss of the neutral marker in the donor strain, under a wide range of parameter values (see **Supplementary Fig. 3**). In fact, the opposite is observed. In all mice long-term maintenance of neutral polymorphism is seen in the donor strain (**Fig. 1B**), even though strong changes in marker frequency occur (**Supplementary Fig. 4**). In striking contrast in the 8 mice where the recipient strain was able to colonize, it lost polymorphism (**Fig. 1B and Supplementary Table 1**). Thus, the probability to maintain polymorphism at a neutral locus within the mouse gut is strain dependent (Binomial Test for proportions, *P* = 0.00028). Under mono-colonization the donor strain also showed a significantly higher abundance than the recipient lineage (Linear mixed-effects model fit by maximum likelihood, *P* = 0.0293) (**Supplementary Fig. 2 and Supplementary Table 1-3**) and its abundance was not significantly different from that under co-colonization.

To determine the evolutionary events that occurred in each strain and quantify their rates of evolution, we performed pool sequencing of clones from each strain after the 88 days of co-colonization (1584 generations, assuming 18 generations per day^14^). A total of 64 mutational events, caused by SNPs (33 events, of which 24 are non-synonymous, 8 are synonymous and 1 is intergenic), IS insertions (10 events), small insertions and deletions (11 events) and large deletions (9 events), were detected in the recipient strain (**Supplementary Table 4**). Five targets of parallel evolution across mice were detected: excision of the cryptic prophage e14, mutations in the repressor of the fructoselysine operon *frlR*, IS insertions in the recombinase *fimE,* in the intergenic region *focA/ycaO* and *mngB/cydA* (**Supplementary Fig. 5A and Supplementary Table 4**). Consistent with being major drivers of gut adaptation in this strain, mutations in most of these targets were found in previous studies^12,14–18^. Across all mice, 9 selective sweeps occurred in the recipient strain, accounting for the loss of polymorphism at the neutral fluorescent marker in most of the mice, except for mouse 7 and 8. In these mice events of HGT need to be invoked to explain the fixation of one of the markers (see below).

The number of mutation events detected in the donor strain was 141. These were caused by SNPs (109 events, of which 65 are non-synonymous, 18 are synonymous and 26 intergenic), IS insertions (8 events), small insertions and deletions (15 events) and large deletions (9 events) (**Supplementary Table 5**). Across 8 mice the ratio of non-synonymous to synonymous mutations (dN/dS) was >1, indicating adaptive evolution. Despite several targets of parallel evolution (**Supplementary Fig. 6A and Supplementary Table 5**), no selective sweeps occurred, consistent with the maintenance of both fluorescence markers in this strain (**Supplementary Fig. 4**). Interestingly, the rate of mutation accumulation, measured as the sum of allele frequencies of the mutations detected over the 1584 generations, was similar between the donor (1.8×10^-3^ (±0.9×10^-3^, S.D.) per generation) and the recipient strain (1.5×10^-3^ (0.3×10^-3^, S.D.) per generation, **Supplementary Fig. 6B**). On one-hand, the donor is a common resident of the mouse gut^11^, and thus could be better adapted to this environment than the recipient, which has been propagated in laboratory environments for many passages. On the other hand, the perturbation of the microbiome composition due to antibiotic treatment could boost its rate of evolution^19^. For the recipient strain a strong signal of metabolic adaptation to the mouse diet was revealed in *in vitro* fitness assays (**Supplementary Table 6**). Evolved clones of this strain showed a large fitness increase when grown in mouse food (**Supplementary Fig. 7 and 8**). In contrast, for the donor strain no clear signal of metabolic adaptation to the mouse diet was found (**Supplementary Fig. 7 and 8**).

A high frequency of HGT events from donor to the recipient *E. coli* lineage marked the evolution of the recipient strain (except in mouse 2) (**Fig. 2A)**. These events were caused by the transfer of two phages (Nef and KingRac) and two plasmids, as previously seen^10^. Strikingly, in one of the mice, mouse 7, a unique transduction event occurred. This involved the transfer of 93712 bp of the genomic region of the donor strain, located adjacent to the prophage KingRac, to the recipient strain (**Fig. 2B)**. This region contains several genes, including an important core gene of *E. coli*: *fnr*. This fumarate and nitrate reductase is an oxygen-responsive transcriptional regulator required for the switch from aerobic to anaerobic metabolism^20^, which is absent in the K12 strain (our ancestral recipient), possibly due to its history of propagation across labs. Our results however show that *fnr* (and several other genes) can be regained by HGT in the gut *via* an evolutionary event consistent with lateral transduction^21^ by prophage KingRac. Both Illumina and Nanopore sequencing (**Supplementary Table 4**), and PCR typing (**Fig. 2A**) confirm this remarkable event, whose frequency steadily increases along the colonization in this mouse (**Supplementary Fig. 5B**). Lateral transduction has been reported to occurred in laboratory conditions^21^ and inferred from comparative genomics^22^. Here we show that it can be caught in real time when two strains of *E. coli* evolve in the mammalian intestine.

**Fig. 2:**
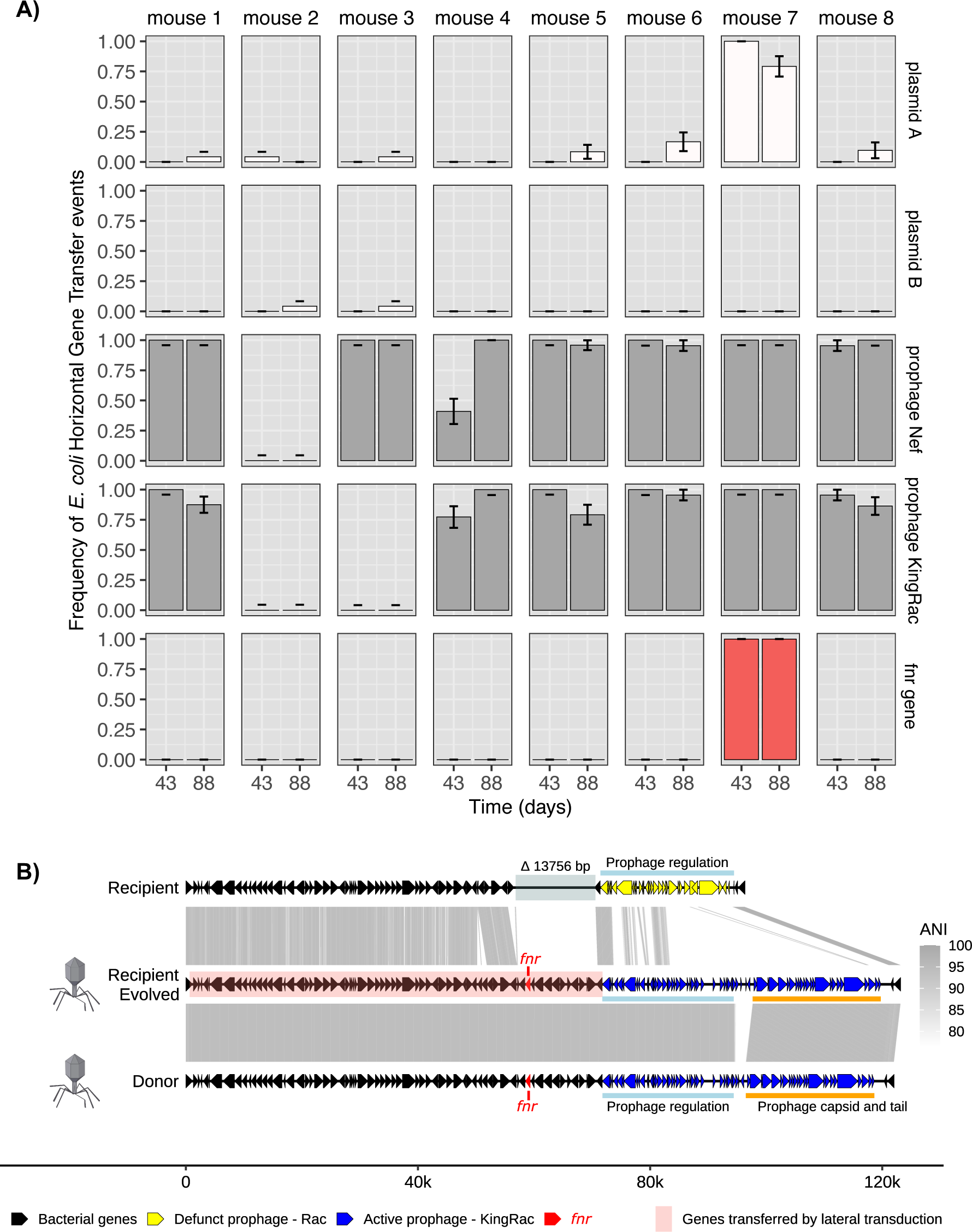
Evolutionary dynamics of HGT from donor to recipient *E. coli* strains. **(A)** Frequency of *E. coli* HGT events from donor to recipient strain at days 43 and 88 after colonization. Error bars represent the standard error of the mean (S.E.M). **(B)** Schematic representation of the massive number of genes gained by the recipient strain, likely by lateral transduction *via* the KingRac prophage. Within the genes gained is *fnr*, a core gene present in the majority of *E.coli* sequenced isolates. Gene arrow representation is to scale.

**Supplementary Information** includes Supplementary Tables.

## Acknowledgments

We would also like to thank the personnel of the IGC’s Rodent Facility, Genomic Facility and the Bioinformatics Unit for their assistance as well as Roberto Balbontín for the marked resident clone. N.F. was supported by “Fundação para a Ciência e Tecnologia” (FCT), fellowship SFRH/BPD/11075/2015, H.C.B. by a cooperation agreement between IGC and the University of Cologne. This work was also supported by project by national funds from FCT and from ERC-2022-ADG 101096203 EvoInHi, funded by the European Union. Views and opinions expressed are however those of the authors only and do not necessarily reflect those of the European Union or ERC. Neither the European Union nor the granting authority can be held responsible for them.

## Author contributions

I.G. designed and coordinated the study. N.F., E.S. performed the experiments and H.C.B. performed the bioinformatic analysis. D.G. performed the PCR typing and M.M. performed the simulations. I.G., N.F., E.S. and H.C.B. analysed the results and wrote the manuscript.

## Author Information

The authors have declared no competing interests. Correspondence and requests for materials should be addressed to I.G.

## Data and materials access

Genome sequencing data has been deposited with links to BioProject accession number PRJNA1019258 in the NCBI BioProject database (https://www.ncbi.nlm.nih.gov/bioproject/). The code for the evolutionary model of accumulation of beneficial mutations is available on the GitHub platform at https://github.com/isabelgordo/HGTransferUnderStrainCoexistence.

---

**Supplementary Fig. 1:**
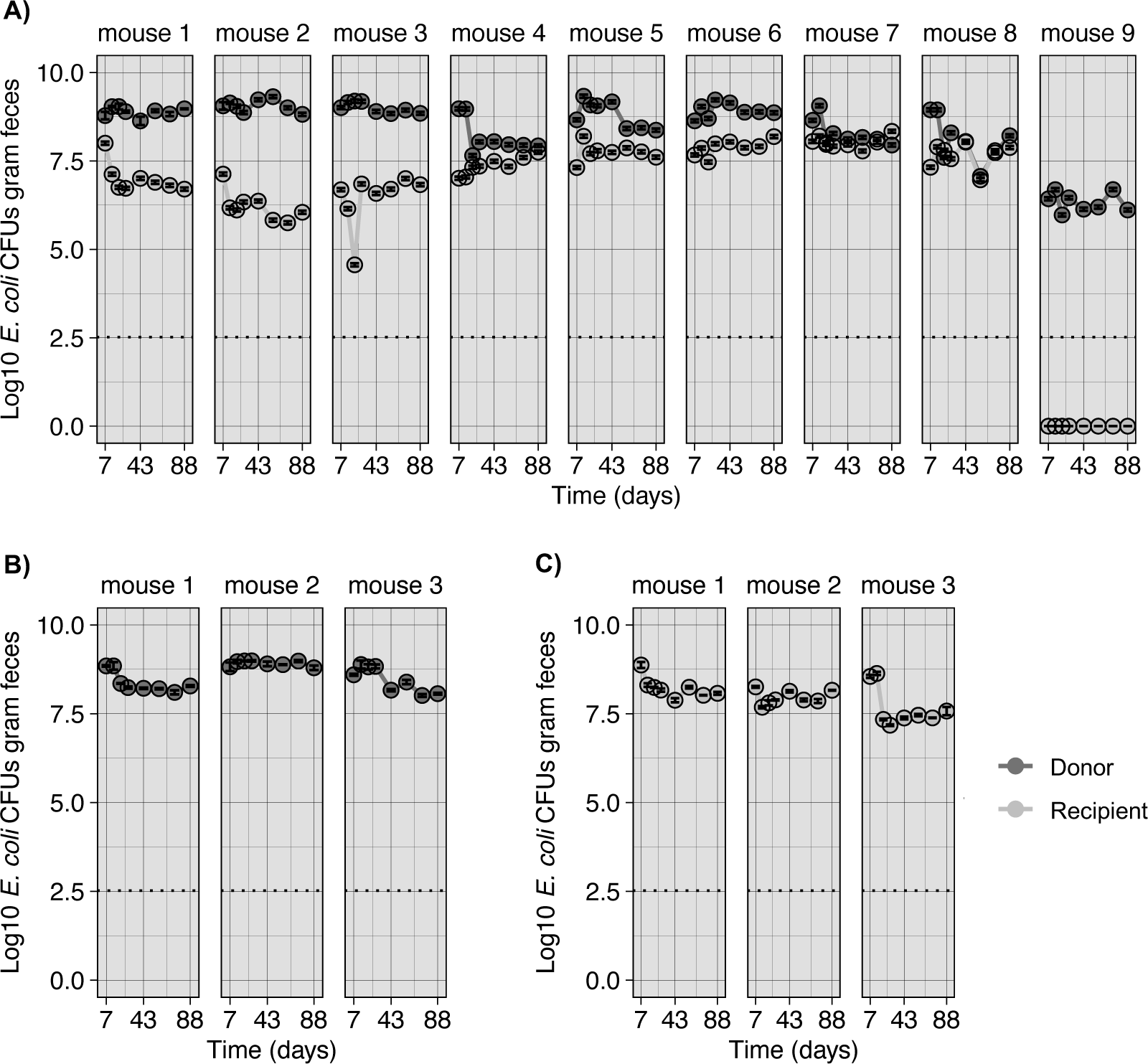
Mean load of the donor and recipient *E. coli* loads across time in each mouse gut. **(A)** Gut bicolonization with donor and recipient strains. **(B)** Gut monocolonization with the donor strain. **(C)** Gut monocolonization with the recipient strain. Error bars represent the Standard Error (2*SE).

**Supplementary Fig. 2:**
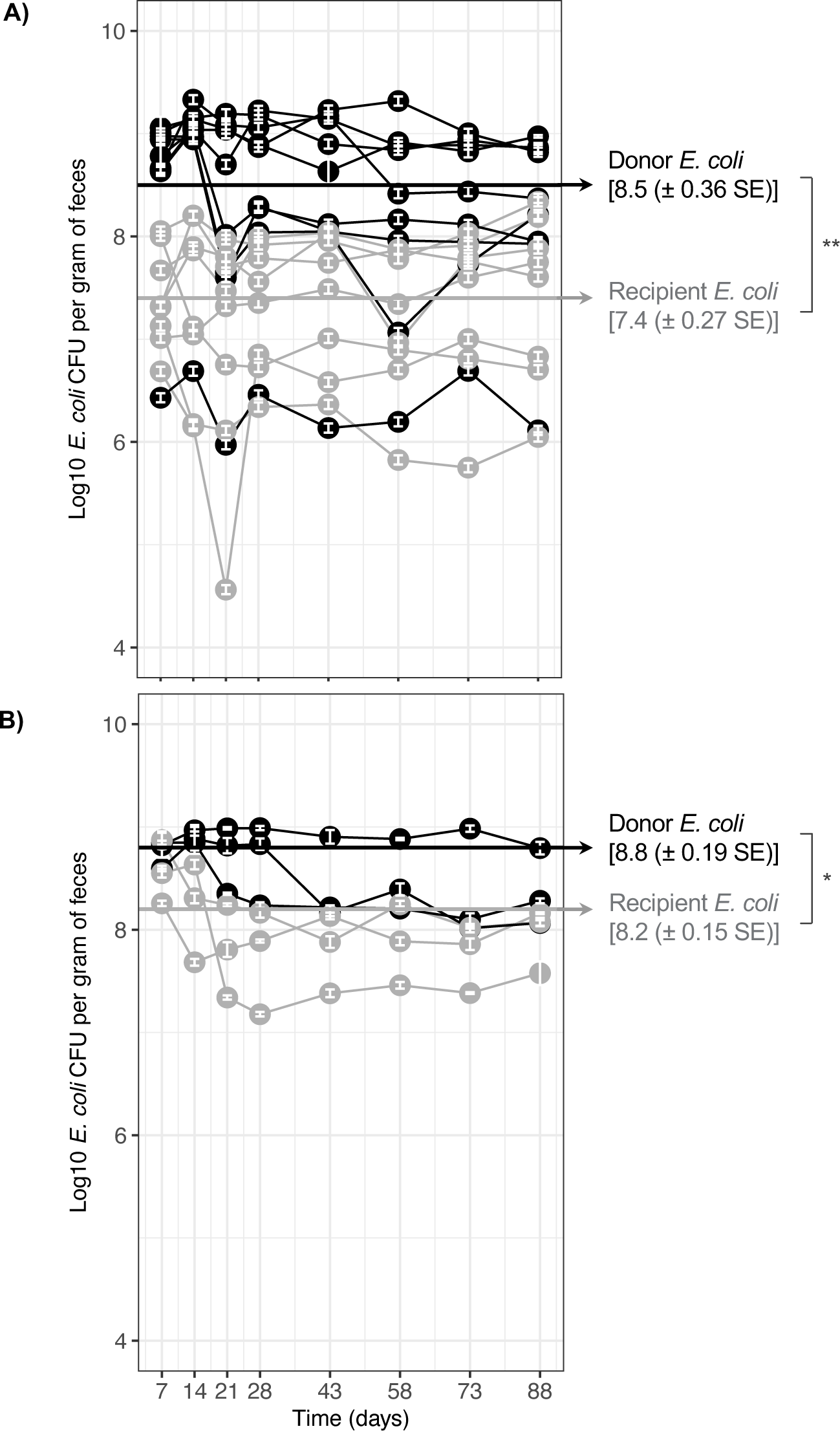
Density of the *E. coli* strains along time in the mouse gut. **(A)** Donor and recipient strains colonizing together: the loads across time of donor and recipient *E. coli* strains are significantly different (Linear mixed-effects model fit by maximum likelihood, p = 0.0074). **(B)** Donor and recipient strains colonizing alone: the loads across time of resident and invader *E. coli* strains are significantly different (Linear mixed-effects model fit by maximum likelihood, p=0.0293). Error bars represent the standard error of the mean (2*S.E.M.).

**Supplementary Fig. 3:**
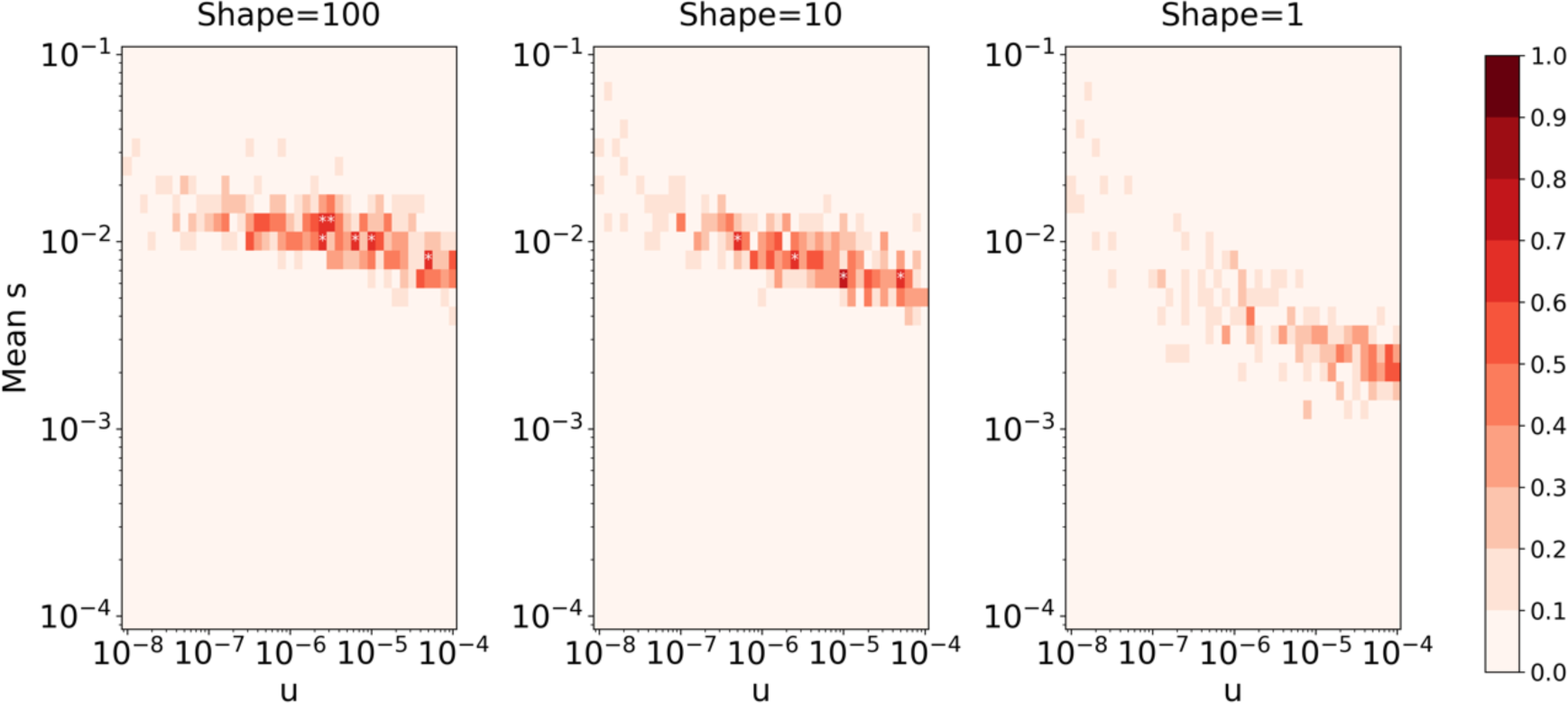
Prediction of simulations of directional selection. The heatmap shows the proportion of simulations compatible with experimental data of fluorescent marker abundances of each strain for each set of parameters (u and mean s). Data-compatibility is defined as maintaining only one marker in the invader strain and the two markers in the resident strain. Parameters were chosen as follows: total population N=10^6^, initial proportion of invaders p=0.1, number of generation g=1600 (88 days with 18 generation per day as in the experiment). The mutation rate u was tested across a large range of values, as well as the mean selective effect s of newly arising beneficial mutations. Mutations were assumed to follow a gamma distribution and three different shapes of this distribution were considered: 100, which gives similar results as those in a model that assumes that all mutations have the same *s* value (left heatmap); 10 (middle heatmap) and 1, which corresponds to an exponential distribution (right heatmap). 10 replicates were simulated for each parameter set. A white star indicates that a significant number of them was compatible with the experimental results of the 8 mice (Fisher exact test). Globally, these simulations show that these simple models could only explain the observed data under very restrictive sets of parameters (mainly on the mean s of the selective effect distribution and on its variance, which has to be very small).

**Supplementary Fig. 4:**
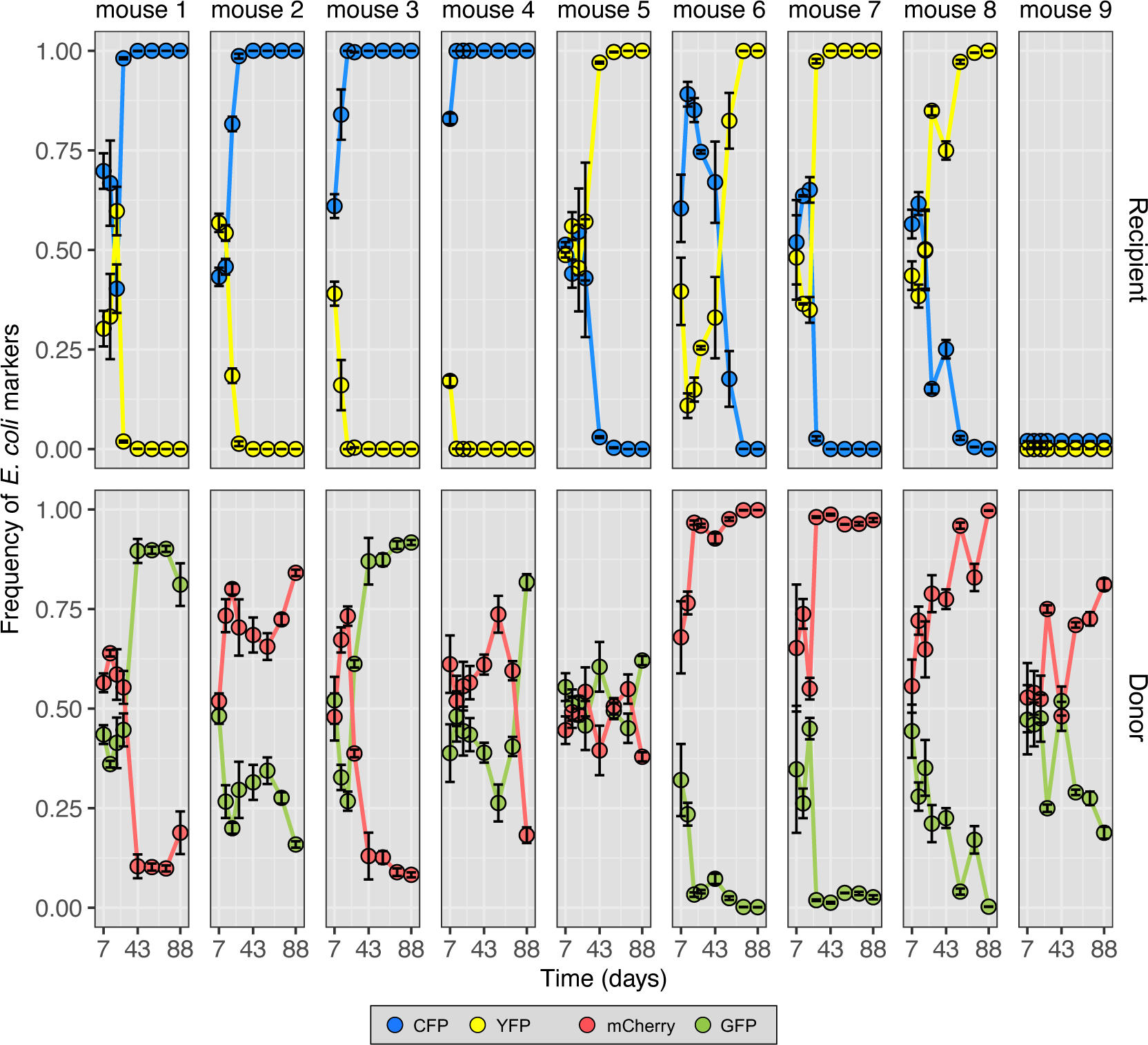
Frequency dynamics of the neutral fluorescent markers of each strain. Frequency of the fluorescent markers in the recipient and donor *E. coli* lineages across ∼3 months (88 days) in different mice (n = 9). Error bars represent the standard error of the mean (2*S.E.M.).

**Supplementary Fig. 5:**
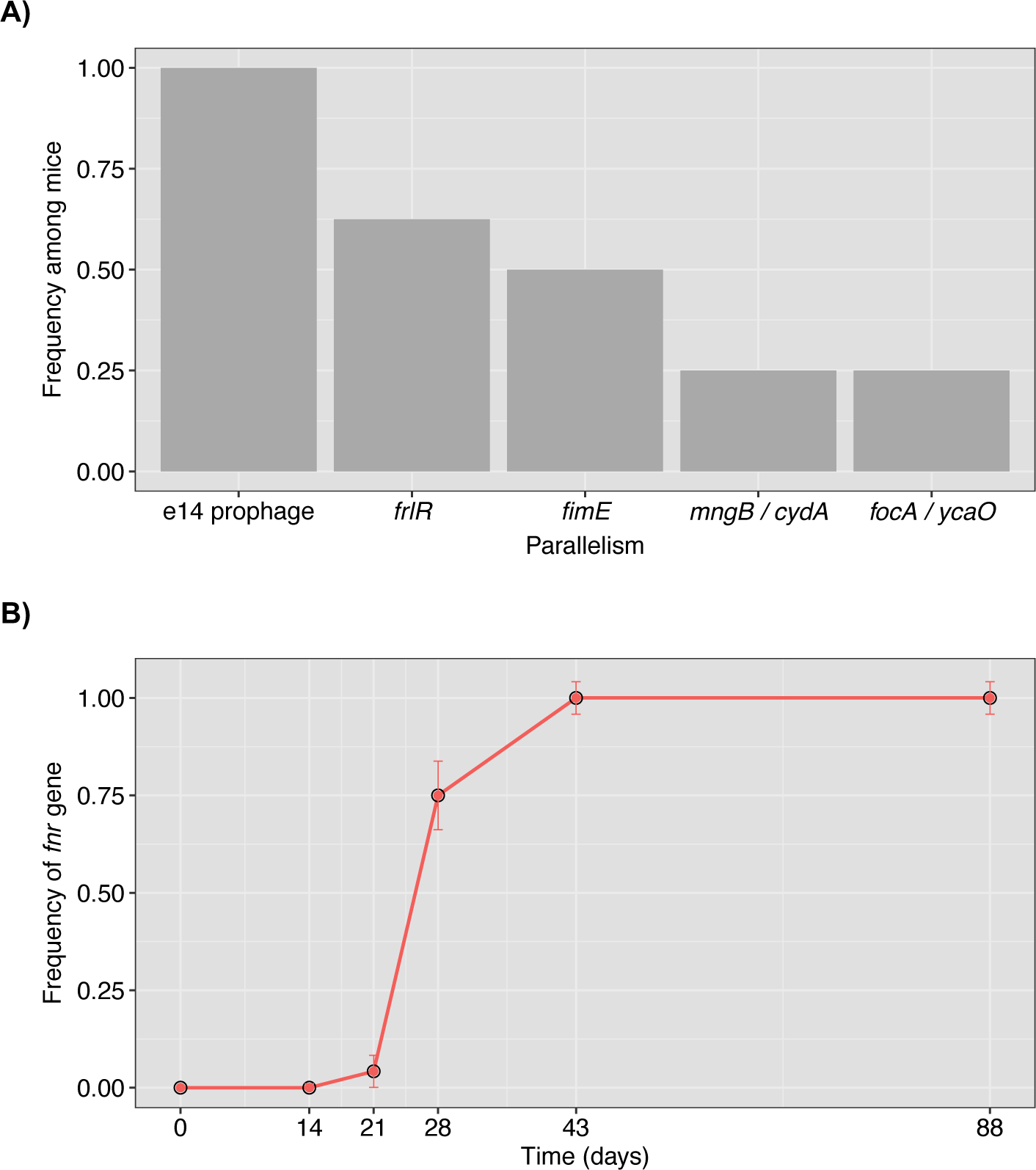
Major targets of adaptation in the recipient strain identified *via* parallel evolution and a HGT event causing a massive transfer of genes. **A)** Frequency of mice (out of 8), where the genetic target was found mutated (only targets mutated in more than one mice are considered adaptive). **B)** Dynamics of the spread of evolved clones which acquired a large lateral gene transfer event (which includes the *fnr* gene, typed here by colony PCR) located next to the prophage KingRac.

**Supplementary Fig. 6:**
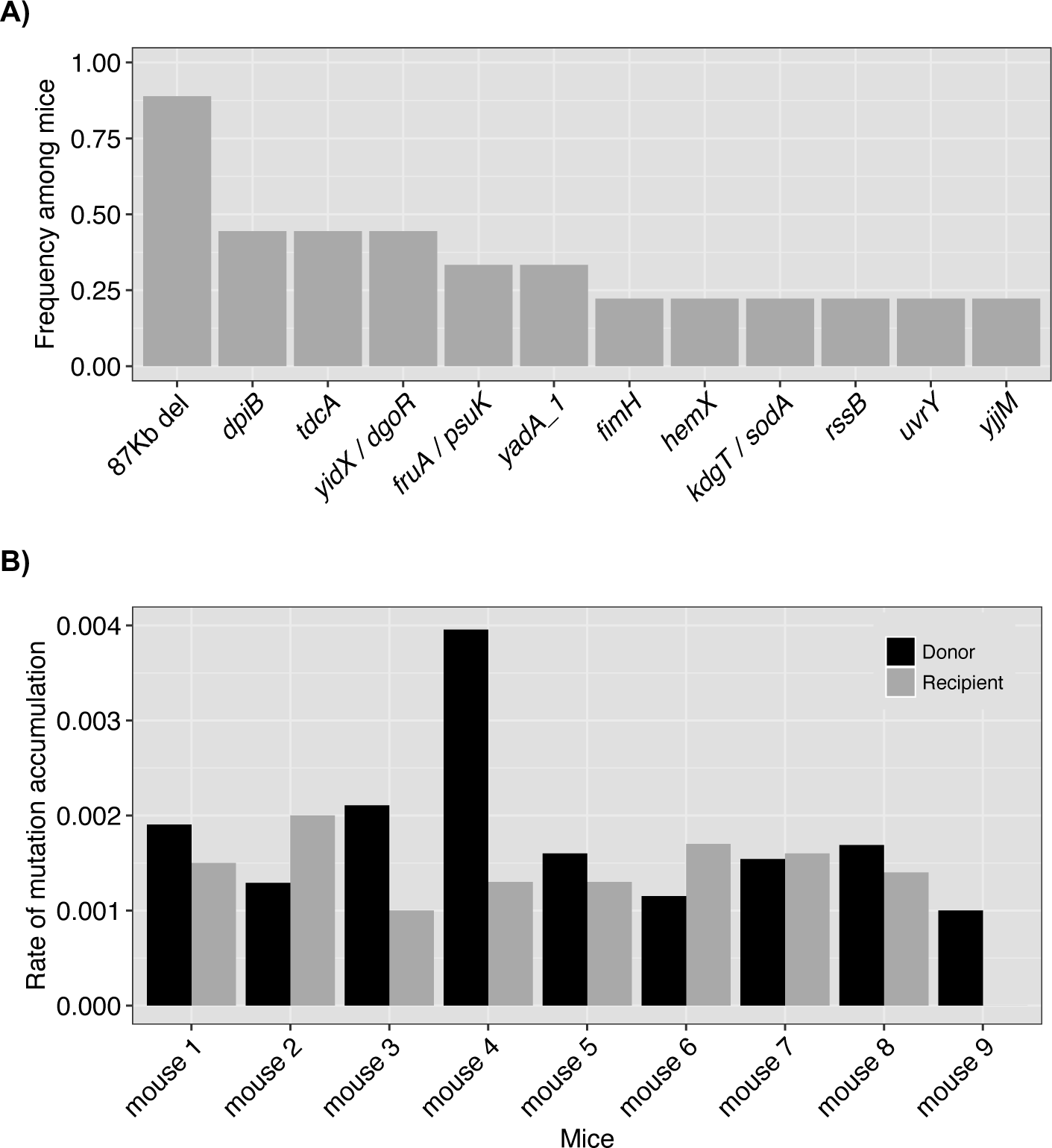
**A)** Targets of parallel evolution identified in the donor strain. Y-axis shows the frequency of mice (out of 9) where genetic target was found mutated. **B)** Rates of mutation accumulation of each strain (donor in dark grey, recipient in light grey) in each mouse. Rates were calculated by summing the allele frequencies of the evolutionary events detected in each strain after 88 days of colonization of the mouse gut. In mouse 4 the donor strain acquired a mutation in *uvrY* that reached a frequency of 44%, which may have caused an increase in mutation rate, leading to a higher rate of mutation accumulation. In mouse 9 the recipient strain failed to achieve persistent colonization.

**Supplementary Fig. 7:**
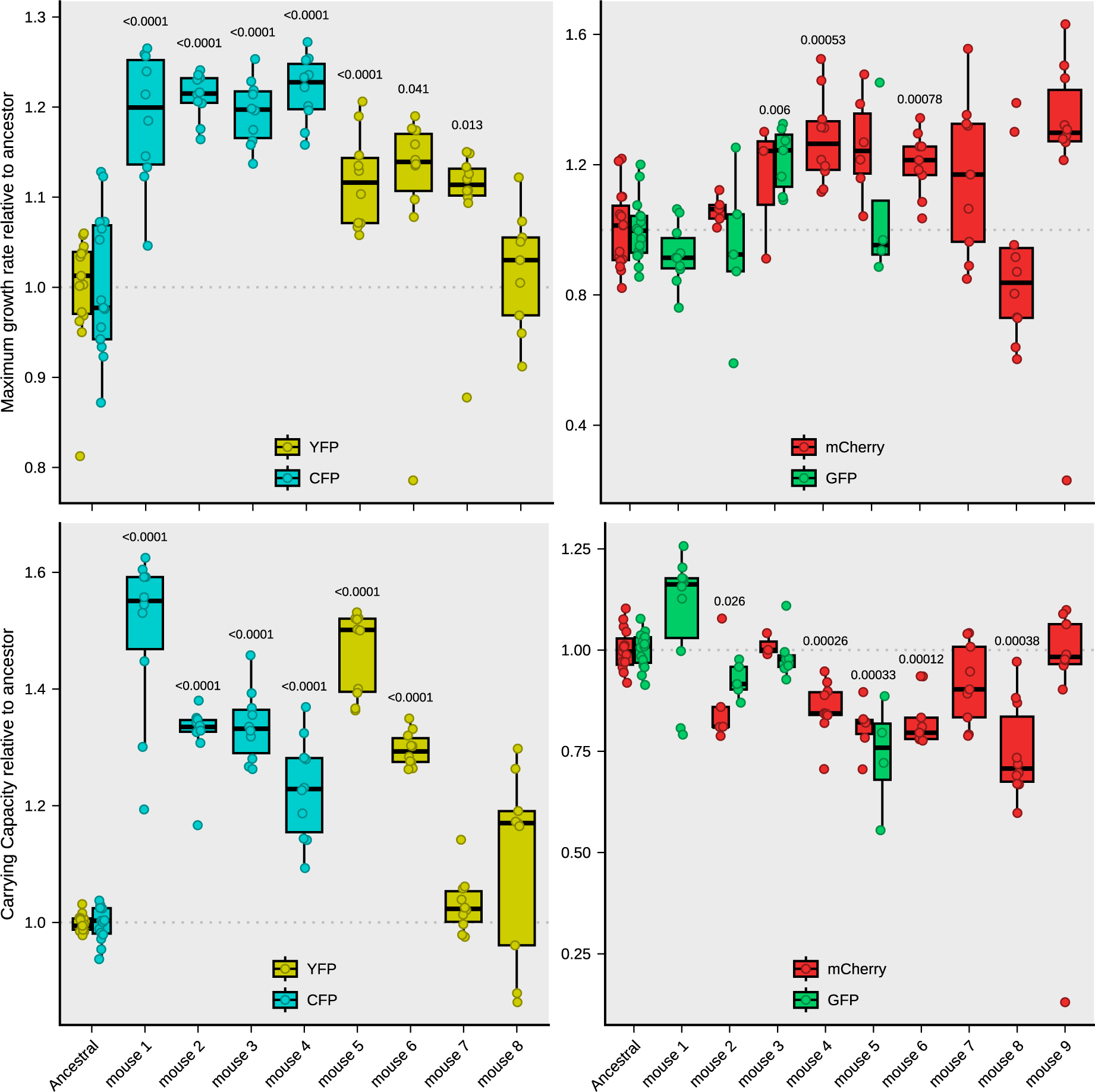
Fitness assays *in vitro* of evolved clones when grown in mouse food. **A)** Relative maximum growth rate (μ/μ_anc_) of clones of the recipient strain sampled after 3 months of gut colonization of each mouse. The values of μ_anc_ (ancestor clones of the recipient strain) are 0.27 h^-1^ (S.D. = 0.02) for the CFP-labeled clone and 0.28 h^-1^ (S.D. = 0.02) for the YFP-labeled clone when grown in mouse food. **B)** Relative maximum growth rate (μ/μ_anc_) of clones of the donor strain. The values of μ_anc_ for the donor strain in mouse food are 1.08 h^-1^ (S.D. = 0.10) for the sfGFP-labeled clone and 1.11 h^-1^ (S.D. = 0.13) for the mCherry-labeled clone. **C)** Relative maximum carrying capacity of clones of the recipient strain sampled after 3 months of gut colonization of each mouse. **D)** Relative maximum carrying capacity of clones of the donor strain sampled after 3 months of gut colonization of each mouse.

**Supplementary Fig. 8:**
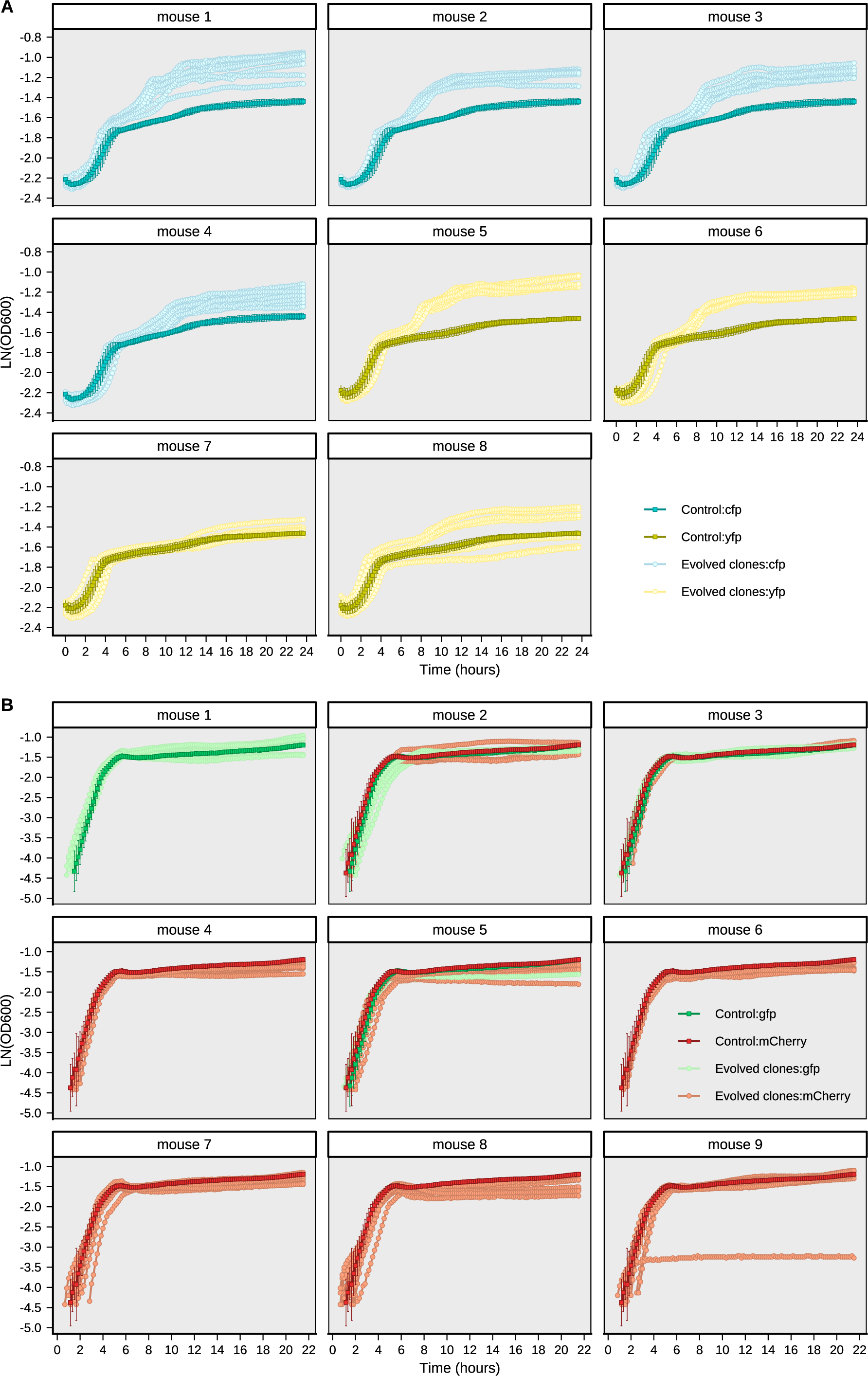
Growth curves of ancestors and evolved clones in mouse food. **A)** Growth curves of ancestor and evolved clones of the recipient strain sampled after 3 months of evolution. **B)** Growth curves of ancestor and evolved clones of the donor strain sampled after 3 months of evolution.

## Methods

### Escherichia coli strains

To facilitate the isolation from the mouse feces all the *E. coli* strains used in this study express fluorescent proteins and carry antibiotic resistance markers. The recipient *E. coli* strains express either a Yellow or a Cyan Fluorescent Protein, namely (YFP) or (CFP), respectively. Each one also carries a streptomycin resistance marker, as well as ampicillin (YFP) or chloramphenicol (CFP) resistance markers. Regarding the resident donor *E. coli* strains, these express either a Red (mCherry) or a Green Fluorescent Protein (sfGFP) and a chloramphenicol resistance marker. *E. coli* clones were grown at 37°C under aeration in liquid media Lysogeny Broth (LB) from SIGMA — or lactose supplemented McConkey agar plates and LB agar plates. Media were supplemented with antibiotics streptomycin (100 µg/mL), ampicillin (100 µg/mL) or chloramphenicol (30 µg/mL) when specified. Serial plating of 1X PBS dilutions of feces in LB agar plates supplemented with the appropriate antibiotics were incubated overnight and YFP, CFP, mCherry and sfGFP-labeled bacterial numbers were assessed by counting the fluorescent colonies using a fluorescent stereoscope (Zeiss Stereo Lumar V12). The detection limit for bacterial plating was ∼300 CFU/g of feces^10^.

### *In vivo* evolution experiment

We used a mouse gut colonization model where mice (*Mus musculus*) were supplied by the Rodent Facility at Instituto Gulbenkian de Ciência (IGC). Animals were kept co-housed to homogenize the mouse microbiota during treatment with streptomycin (5 g/L) in the drinking water for 7 days and the following 2 days, streptomycin treatment was absent to clean the gut from antibiotic traces^23^. Afterwards the animals were inoculated by gavage with 100 µL of an *E. coli* bacterial suspension of ∼10^8^ colony forming units (CFUs) and housed individually with drinking water not supplemented with streptomycin. Six- to eight-week-old C57BL/6J non-littermate female mice (n=15) were kept in individually ventilated cages under specified pathogen free (SPF) barrier conditions at the IGC animal facility. Nine mice were colonized by intragastrical gavage with two strains of *E. coli* (the recipient and the donor strains). Three with the recipient strain and another 3 with the donor strain. Fecal pellets were collected during ∼3 months (88 days) and stored in 15% glycerol at -80°C for later analysis. This research project was ethically reviewed and approved by the Ethics Committee of the Instituto Gulbenkian de Ciência (license reference: A009.2018), and by the Portuguese National Entity that regulates the use of laboratory animals (DGAV - Direção Geral de Alimentação e Veterinária (license reference: 008958). All experiments conducted on animals followed the Portuguese (Decreto-Lei n° 113/2013) and European (Directive 2010/63/EU) legislations, concerning housing, husbandry and animal welfare.

### Whole-genome sequencing and analysis pipeline

DNA was extracted with Phenol-Chloroform^24^ from *E. coli* populations (mixture of >1000 clones) or a single clone growing in LB plates supplemented with antibiotic to avoid contamination. DNA concentration and purity were quantified using Qubit and NanoDrop, respectively. The DNA library construction and sequencing were carried out by the IGC genomics facility using the Illumina Nextseq2000 platform. Processing of raw reads and variants analysis was based on previous work^12,18^. Briefly, sequencing adapters were removed using fastp^25^ and raw reads were trimmed bidirectionally by 4bp window sizes across which an average base quality of 20 was required to be retained. Further retention of reads required a minimum length of 100 bps per read containing at least 50% base pairs with phred scores at or above 20. BBsplit^26^ was used to remove likely contaminating reads. Separate reference genomes were used for the alignment of recipient (K-12 substrain MG1655; Accession Number: NC_000913.2) and donor (Accession Number: SAMN15163749) *E. coli* genomes. Annotation of IS elements in the donor genome was performed using ISEScan^27^ (version 1.7.2.3), and only complete IS were considered for the annotation of the donor reference genome. As mouse 8 donor and recipient samples were contaminated with reads from *Enterobacter hormachei*, BBSplit was run including genomes from *E. hormachei* (Accession Numbers: CP104691.1, CP104689.1, CP056649.1, CP044335.1, CP051132.1, CP116960.1, and CP019889.1). The donor sample from mouse 4 was contaminated with reads from the recipient, in this sample BBSplit was run including the reference genome of the recipient (K-12 substrain MG1655; Accession Number: NC_000913.2). Mutations were identified using the 0.37.1 version of the BRESEQ pipeline^28^ using the reference genome for the recipient (K-12 substrain MG1655; Accession Number: NC_000913.2) and the IS annotated donor genome (based on Accession Number: SAMN15163749) *E. coli* genomes, with polymorphism option on and default settings except for a) polymorphism minimum variant coverage of 5 reads; b) base quality cutoff of 30; c) minimum mapping quality of 20. Final average alignment depths ranged from 96 to 234 for the recipient populations (median = 200), and from 100 to 466 for the donor populations (median = 308). The ancestral strains of the donor and recipient were sequenced and analysed as described for the populations. BRESEQ output from the populations was then conservatively filtered using a custom R script with the following steps: a) removal of the predicted mutations already present in the ancestral strains ran with the same parameters as the sequenced populations; b) BLAST all the reads of each predicted mutations against the recipient and donor reference genomes (if reads that 100% match with the incorrect reference genome are at a frequency above 0.01, the predicted mutation is removed from the list and considered as a potential HGT event); c) removal of predicted mutations if they are detected in the reads of the ancestral clones at a frequency higher than 0.015 and with more than 3 reads. All the remaining putative variants were verified manually in IGV^29^.

### Nanopore sequencing

DNA concentration and purity were accessed as described above, and DNA library construction and sequencing were carried out by the IGC genomics facility using the Oxford Nanopore Tecnologies (ONT) and MinION. *De novo* assembly was performed using the Flye^30^ pipeline (version 2.9.2) with the following parameters: a) ONT high-quality reads; b) estimated genome size of 4.7m; c) read error rate of 0.05. The assembly was then annotated with Prokka^31^ using the annotated proteins from the reference genome of the recipient (K-12 substrain MG1655; Accession Number: NC_000913.2) as trusted proteins to first annotate from. Putative prophage regions were previously identified^12^. Mutations were identified using the 0.37.1 version of the BRESEQ pipeline^28^ using the reference genome for the recipient (K-12 substrain MG1655; Accession Number: NC_000913.2) with default settings. Average Nucleotide Identity (ANI) between the recipient, donor, and evolved recipient strains for the genomic region located adjacent to the prophage KingRac was calculated using fastANI^32^ (version 1.33) with default parameters except: a) fragLen = 100.

### PCR detection of ∼69Kb (*repA*) and ∼109Kb (*repB*) plasmids

Specific primers for the amplification of *repA* and *repB* genes, were used to determine the frequency of the 68935 bp (∼69 Kb) and 108557 bp (∼109 Kb) plasmids, respectively, in the invader *E. coli* populations at days 43 and 88 after gut colonization.

The primers used for *repA* gene were:

repA-Forward: 5’-CAGTCCCCTAAAGAATCGCCCC-3’ and repA-Reverse: 5’-TGACCAGGAGCGGCACAATCGC-3’.

For *repB* the primer sequences were:

repB-Forward: 5’-GTGGATAAGTCGTCCGGTGAGC-3’ and repB-Reverse: 5’-GTTCAAACAGGCGGGGATCGGC3’.

PCR amplification of plasmid-specific genes was performed in randomly isolated clones from the invader evolved *E. coli* populations. PCR reactions were performed in a total volume of 25 μL, containing 1 μL of each clone growth in liquid LB media, 1X Taq polymerase buffer, 200 μM dNTPs, 0.2 μM of each primer and 1.25 U Taq polymerase. PCR reaction conditions: 95°C for 3 min, followed by 35 cycles of 95°C for 30 s, 65°C for 30 s and 72°C for 30 s, finalizing with 5 min at 72°C. DNA was visualized on a 2% agarose gel stained with GelRed and run at 160 V for 60 min.

### PCR detection of the Nef and KingRac prophages

PCR amplification of phage-specific genes was performed as previously^11^ to determine the frequency of lysogens in the recipient *E. coli* population at days 43 and 88 after gut colonization.

### PCR detection of the *fnr* gene

Specific primers for the amplification of *fnr* gene, were used to determine its frequency in the recipient *E. coli* populations at days 43 and 88 after gut colonization. The primers used were: *fnr*-Forward: 5’-CATTTAGCTGGCGACCTGGTGG-3’, *ynaJ*_fw: 5’-TTCAGAGCAGACAACGGTGA-3’ and *ttcA*_rv: 5’-GGGCGTATCGAGACGATGTT-3’. PCR amplification of the *fnr* gene was performed in randomly isolated clones from the recipient evolved *E. coli* populations. PCR reactions were performed in a total volume of 25 μL, containing 1 μL of each clone grown in liquid LB media, 1X Taq polymerase buffer, 200 μM dNTPs, 0.2 μM of each primer and 1.25 U Taq polymerase. PCR reaction conditions: 95°C for 3 min, followed by 35 cycles of 95°C for 30 s, 60°C for 30s and 72°C for 1 min, finalizing with 5 min at 72°C. DNA was visualized on 1% agarose gel stained with Xpert Green.

### Growth assays *in vitro*

A pre-culture was prepared by inoculating the previously isolated single clones from the frozen stock to 150 μl MM + glucose [0.5 mM] in a 96-well plate, followed by incubation for 24 h at 37°C with 600 rpm in a benchtop shaker (Thermo-Shaker PHMP-4, Grant). To generate single clone growth curves the wells of a BioscreenC Honeycomb plate (Thermo Scientific) were filled with 295 μl culture medium and inoculated with 5 μl of pre-culture diluted in PBS, aiming at 10^5^ cells. The OD600 was measured with the BioscreenC, (Oy Growth Curves Ab Ltd.) for 24 h at 37°C with reading intervals every 10 minutes under continuous shaking with medium amplitude. The culture medium from mouse food was prepared using autoclaved Rat and Mouse food No.3 (SDS Special Diets Service). 75 g pellets of mouse food per liter were dissolved in sterile ddH_2_O, then centrifuged and the supernatant filtered through a 0.22 μm PES membrane (Avantor, VWR). For each clone a maximum growth rate was estimated from 5 measurements of optical density (OD) during the exponential phase of the growth, by calculating the slope of Ln(OD – OD_blank_) in a linear regression, which led to the highest growth rate and a R^2^>0.9.

### Simulations of directional selection

A simple evolutionary model of accumulation of beneficial mutations was simulated to enquire under which conditions the data obtained from measuring the abundance of neutral markers: two resident strains maintained, one invading strain eliminated and the other maintained. Beneficial mutations were assumed to be Poisson distributed with a constant mutation rate (u) and the total population size of both strains was constant (*N*=10^6^). The model assumes non-overlapping generations, and each new generation is drawn *via* multinomial sampling of the precedent one, with probabilities proportional to the relative fitness of a mutant strain. The selective advantage of each new given by a new mutation is assumed to be fixed or drawn from an exponential distribution.

The total population is divided into 4 types (modeling the different fluorescence in the experiment, each with an initial equal fitness (set to 1), and the initial population sizes of each type were similar to those in the experiment: (0.05,0.05,0.45,0.45). 10 replicas for each parameter set were done. A large range of parameters was explored: selective advantage from 10^-4^ to 10^-1^ and mutation rate from 10^-8^ to 10^-4^, both with increments on a log scale. A Fisher’s exact test was used to assess whether the difference between the simulated and the experimental results ones were not significantly different.

### Statistical analysis

A Linear mixed model (R package nlme, v3.1^33^) was used to analyze the load temporal dynamics of the invader and resident *E. coli* lineages, while colonizing the mouse gut. A Binomial Test was used to compare strain polymorphism proportions. T-tests were used to compare differences in maximum growth rate between ancestor and evolved clones, and a paired T-test was used to compare rates of mutation accumulation of each strain. A P-value < 0.05 was considered for statistical significance.

